# Loss of transcriptional factor *Zbtb33* fails to induce clonal hematopoiesis in mice but plays a role in tumor immunity

**DOI:** 10.1101/2025.04.10.648125

**Authors:** Yuhang Li, Kaiping Luo, Jingjing Liu, Ge Dong, Zhigang Zhao, Zhigang Cai

**Affiliations:** Department of Oncology, Tianjin Medical University Cancer Institute and Hospital, National Clinical Research Center for Cancer, Key Laboratory of Cancer Prevention and Therapy, Tianjin’s Clinical Research Center for Cancer, Tianjin, China; State Key Laboratory of Experimental Hematology, Tianjin Medical University, Tianjin, China; Tianjin Key Laboratory of Inflammatory Biology, Department of Pharmacology, School of Basic Medical Science, Tianjin Medical University, Tianjin, China; The Province and Ministry Co-sponsored Collaborative Innovation Center for Medical Epigenetics, School of Basic Medical Science, Tianjin Medical University, Tianjin, China; Department of Medical Oncology, Tianjin First Central Hospital, School of Medicine, Nankai University, Tianjin, China

**Keywords:** Clonal hematopoiesis, *Zbtb33*, *Tet2*, *Tp53*, Inflammation, Immune regulation

## Abstract

Clonal hematopoiesis (CH) is an early indicator of hematologic malignancies, driven by mutations in hematopoietic stem cells (HSCs) such as TET2 or TP53. Mutations in *ZBTB33* have been implicated in MDS and suggested as a potential driver of CH. However, the role of *ZBTB33* in hematopoiesis and its involvement in CH remains unclear. We generated a *Zbtb33*-knockout mouse strain to elucidate its role in hematopoiesis and the immune system. Our findings indicate that hematopoiesis in *Zbtb33*-defecient mice appeared grossly normal, and competitive bone marrow transplantation assays demonstrated that loss of *Zbtb33* in HSCs did not confer expansional advantage. Introducing the *Zbtb33* mutation into *Tet2*- or *Tp53*-mutation background yielded no synergistical effects. Tumor challenging assays suggested that *Zbtb33* influences cancer immunity response, rather than directly driving CH or myeloid malignancies. In summary, *ZBTB33* deficiency was insufficient to induce clonal hematopoiesis but may have a regulatory role in tumor microenvironment.

**Statement of significance:** Clonal hematopoiesis (CH) is linked to mutations in hematopoietic stem cells, but the role of Zbtb33 in CH remains unclear. To investigate this, we examined the function of Zbtb33 under physiological conditions and in response to external stimuli. Additionally, we explored whether Zbtb33 mutations cooperate with other genetic mutations to drive clonal hematopoiesis.

**Key points:**

1. Loss of Zbtb33 fails to induce clonal hematopoiesis and does not synergize with *Tet* or *Tp53* mutations.
2. However, it plays a significant role in regulating cancer immunity and the tumor microenvironment.

## Introduction

Clonal hematopoiesis (CH) is an age-associated aberrant hematopoiesis phenomena characterized by the presence of certain somatic mutations in hematopoietic stem cells (HSCs), leading to the clonal expansion advantages of these cells and an increase in the proportion of mutant blood cells in circulation.^1,2^ Clonal hematopoiesis is recognized as a pivotal precursor to various hematological malignancies, such as myelodysplastic syndromes (MDS) and acute myeloid leukemia (AML), as well as an independent risk factor for several non-hematological chronic diseases, such as cardiovascular diseases, colitis, and autoimmunity disease. Early detection of CH could potentially allow for interventions that may prevent or delay the onset of associated diseases.^3,4^ Clonal hematopoiesis of indeterminate potential (CHIP) is a significant concept defined by the presence of specific somatic mutations in peripheral blood with variant allele fractions (VAF) exceeding 2%.^4,5^ Among the well-known drivers of CH, genes such as *TET2*, *DNMT3A*, *TP53*, and *JAK2* have been identified in recent clinical and epidemiological studies; and importantly, validated through experimentations using mouse models and/or human cell models, suggesting their causal role in CH.^6,7^

ZBTB33 (Zinc finger and BTB domain containing 33) also known as KAISO, is a member of the BTB/POZ subfamily of zinc finger transcription factors. Aberrant *ZBTB33/KAISO* expression has been observed in multiple hematopoietic malignancies such as chronic myeloid leukemia, chronic myelomonocytic leukemia-like and atypical chronic myelomonocytic-like at the molecular level.^8^ A recent study has also suggested that *ZBTB33* is linked to poor outcomes in acute T lymphocytic leukemia.^20^ Additionally, *ZBTB33* has been implicated in a variety of human malignancies beyond hematopoietic systems, including prostate^11,12^, breast^13,14^, pancreatic^15^, and lung cancers^16^. Its involvement in these cancers indicates a broad impact of *ZBTB33* on tumorigenesis.

The origin (i.e. drivers) and dynamics (i.e. history, trajectory and outcomes) of CH are central to understanding its role in the development of hematological malignancies and other diseases.^17–19^ Our previous studies, corroborated by recent related studies, have suggested that *Tet2* mutations in HSC lead to three distinct outcomes: 1. *Tet2* mutant HSCs have increased self-renew capability; 2. Myeloid-skewing differentiation from the *Tet2* mutant HSCs; 3. *Tet2* mutant mature myeloid cells manifest increased inflammation.^20^ In our most recent study, we demonstrated that DSS-induced colitis cooperate with *Tet2* mutation to promote *Tet2*-defeciency mediated CH.^21^ In the present study, our motivation is to ask if any new CH candidate drivers cooperate with *Tet2* mutation and synergistically promote CH and leukemogenesis.

A study by *Beauchamp et al.* published in *Blood Cancer Discovery* in 2021 has brought to light that *Zbtb33*, emerges as a newly “identified” potential driver of clonal hematopoiesis, using a tailored pipeline to unbiasedly integrate diverse human peripheral blood DNA sequencing datasets from ExAC and TOPMed.^4^ Concurrently, during the course of our study, another research by *Berntein et al.* in *Nature Genetics* in 2024 reported that *Zbtb33* as one of seventeen novel drivers of CH, based on clinical epidemiology only rather than experimental validataions.^22^ Moreover, a mutation pattern in *Zbtb33* has been recurrently detected with high VAFs in multiple clinical trials of MPNs (myeloproliferative neoplasms)/MDS^5–6^, which are inclined towards malignant transformation (i.e. AML).^24,25^ These studies suggest a potential role of *Zbtb33* in clonal hematopoiesis and CH-related hematopoietic malignancies. However, it is important to note that these studies establish “association” rather than “causality”. Thus, the question of whether HSCs carrying *Zbtb33* loss of function mutations enhance self-renewal and expansion advantages remains to be clarified through experiments. Furthermore, there is a clear need for a systematic and *in vivo* investigation into the role of *Zbtb33* in hematopoiesis and in other biological process.

Here, we generated *Zbtb33*-knockout mice (*Zbtb33*-KO) to explore the potential alterations in hematopoiesis under physiological condition or in response to external stimuli. Our experimental results do not support the hypothesis that *Zbtb33/Kaiso* mutations act as drivers of CH; moreover, our results suggest that loss of *Zbt33* does not exhibit synergistic effects with the loss of *Tet2* or *Tp53*. Our preliminary data suggest that *Zbtb33* is with equally abundant expression in both myeloid and T cells, and loss of *Zbtb33* in tumor microenvironment resulted in mitigated tumor growth, suggesting a role for *Zbtb33* in tumor immunity.

## Results

### Generation of Zbtb33 knockout mice and monitoring of Zbtb33^-/^*^y^* (male mutants) *p*eripheral blood

Using CRISPR/Cas9 technology, two sgRNA were designed to target the core region of *Zbtb33* Exon 2 where protein-coding sequences initiate (note *Zbtb33/ZBTB33* is located at X chromosome in mouse and human, **Supplemental Figure 1A**). Recognizing that somatic mutations in *ZBTB33* are predominantly identified in male donors with clonal hematopoiesis^22^, this study focused on male mutants (*Zbtb33^-/y^*). Although high-quality antibodies for ZBTB33 are unavailable, transcriptional depletion of *Zbtb33* was confirmed by bulk RNA-seq of wild-type and mutant bone marrow cells (BMCs) (**Figure 1A**).

**Figure 1.**
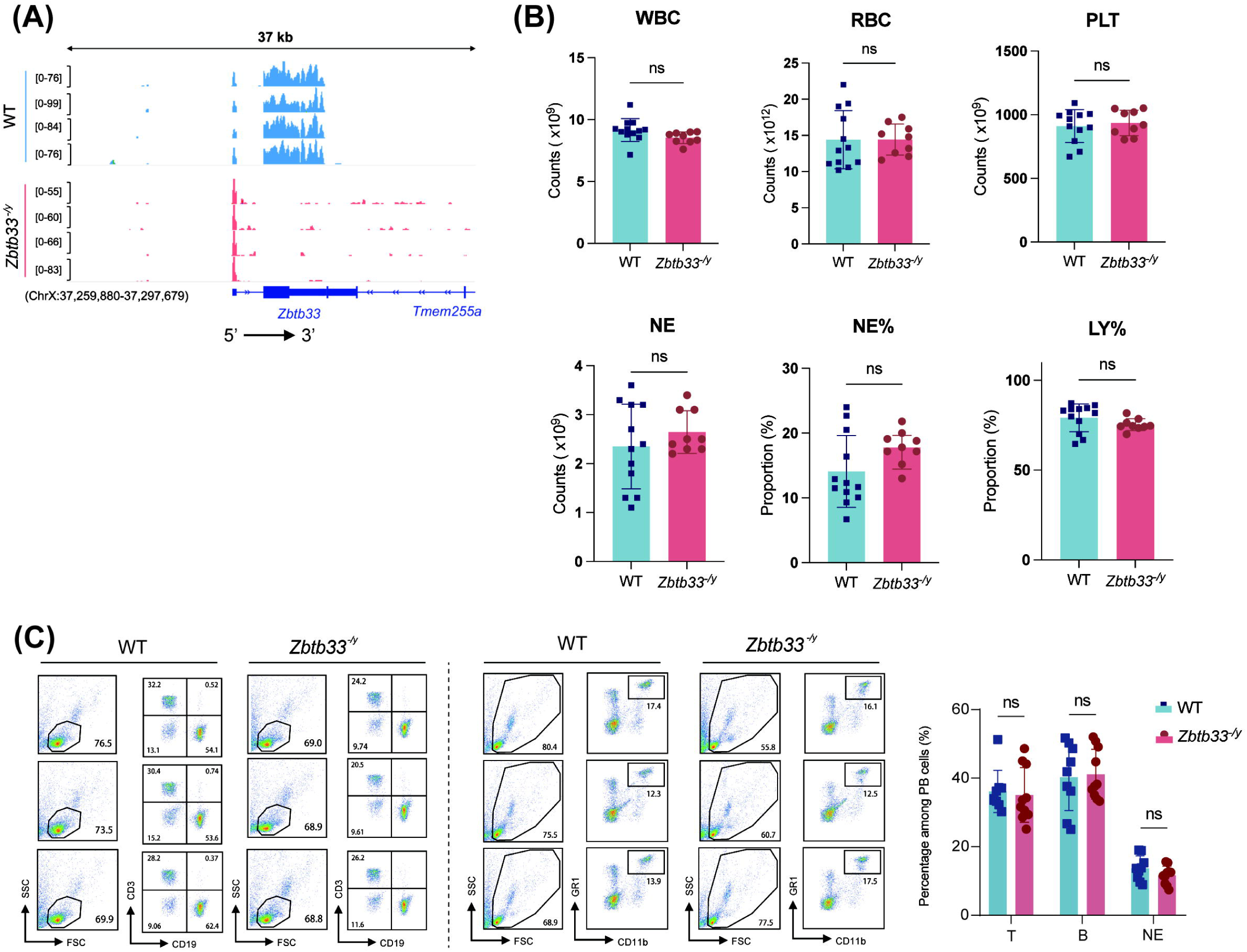
Generation of *Zbtb33* knockout mice and monitoring of *Zbtb33^-/y^* peripheral blood. (**A**) Read alignment using bulk RNA-seq datasets of bone marrow cells from Zbtb33-deficient and wildtype mice. Region for *Zbtb33* locus (Exon-1 and -2) is showing. Exon-1 is preserved but Exon-2 is absent in *Zbtb33^-/y^*, which is consistent to the targeting strategy (see **Supplemental Figure 1A**). (**B**) Peripheral blood (PB) cell counts for adult *Zbtb33* deficient mice and wildtype littermates. (**C**) Representative flow cytometry profiles and quantification of T cells, B cells and Neutrophils in PB. Cell counts and flow cytometry analysis suggest the *Zbtb33*^-/y^ mice manifest normal blood cell compositions.

We performed a comprehensive phenotypic assessment of *Zbtb33^-/y^* male mice. Peripheral blood (PB) parameters were monitored throughout their lifetime and appeared normal compared to controls. At 6-8 months of age, hematological parameters (including WBC, RBC, PLT, NE, NE%, and LY%) showed no significant differences between mutant and wild-type mice (**Figure 1B**). These findings are in concordance with previous findings reported by an independent research group.^26^ Flow cytometry analysis also revealed no significant abnormalities in proportions of T cells (CD19^-^; CD3^+^), B cells (CD19^+^; CD3^-^), or neutrophils (CD11b^+^; GR1^+^) in PB of *Zbtb33^-/y^*mice (**Figure 1C**). These findings were consistent across extended ages (6-12 months), where we confirmed that these parameters remained within the normal ranges (**Supplemental Figure 1B**). In summary, Zbtb33-deficient male mice exhibited no noticeable alterations in PB parameters throughout the examined age range (2-12 months).

### Loss of Zbtb33 is insufficient to affect hematopoietic stem in bone marrow

To investigate the impact of *Zbtb33* deficiency on developmental hierarchy and cellularity of bone marrow, we compared the proportions of hematopoietic stem and progenitor cell (HSPC) subsets between *Zbtb33*-knockout (KO) mice and wild-type (WT) littermates using cytometric analyses (**Figure 2A-F**). No statistically significant differences were observed in the proportions of HSPC subtypes, although a slight, non-significant increase in the proportion of LinLJ cells was noted in *Zbtb33*-KO mice (**Figure 2A**, *P* = 0.09). Similarly, the proportions of LSK (LinLJSca1LJKitLJ) cells and their subpopulations, including hematopoietic stem cells (HSCs, LSK-CD150LJCD48LJ), remained comparable between groups (**Figure 2B-G**). Given the potential role of ZBTB33 in myeloid hematological disorders such as MDS/MPNs, we further examined myeloid hematopoietic progenitor subpopulations. Regrettably, no significant differences were found in the proportions of myeloid progenitors (MPs, Lin^-^Sca1^-^cKit^+^) and common myeloid progenitors (CMPs, Lin^-^Sca1^-^cKit^+^CD16^-^CD34^+^) between *Zbtb33*-knockout mice and WT mice. (**Figure 2H**). However, in terms of the mature cells in bone marrow, the proportion of neutrophils was significantly higher in *Zbtb33*-KO mice, while T cells (CD3LJ) and B cells (CD19LJ) remained unchanged, suggesting selective enhancement of neutrophil differentiation (**Figure 2I, P < 0.05**). Collectively, these findings suggested that the loss of *Zbtb33* was insufficient to affect the proliferation and expansion of HSPC but moderately enhance cell differentiation and maturation of certain myeloid lineage (neutrophil) in bone marrow.

**Figure 2.**
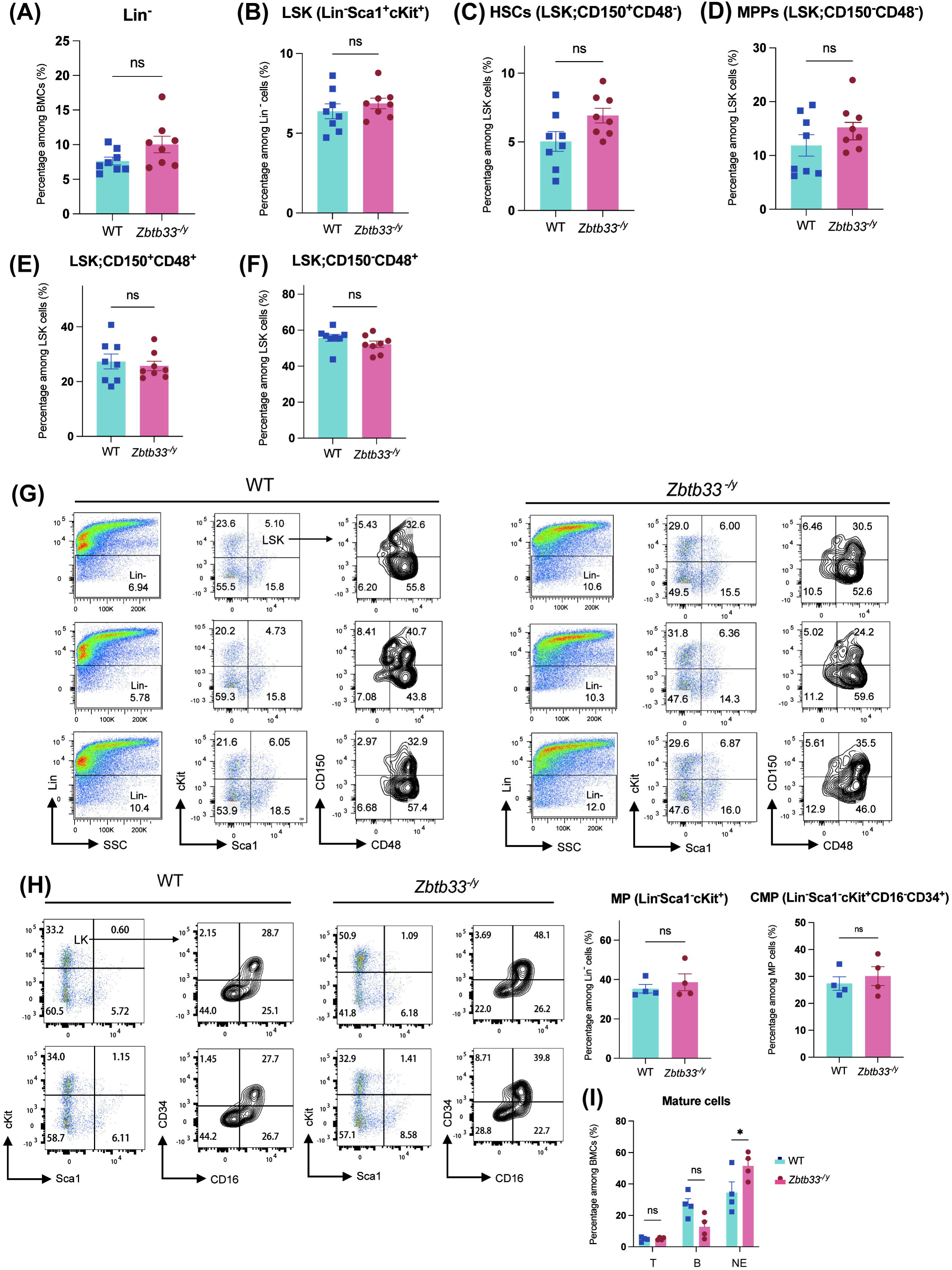
Loss of *Zbtb33* subtly affects hematopoietic stem and progenitor cells in bone marrow. **(A)** Percentage of Lin^-^ cells in the bone marrow of *Zbtb33* knockout mice compared with the wildtype littermates. **(B)** Percentage of Lin^-^Sca1^+^cKit^+^ cells (LSK) in Lin^-^ cells in the BM. (**C**-**F**) Percentage of hematopoietic stem cells (Lin^-^Sca1^+^cKit^+^CD150^+^CD48^-^, HSCs) (**C**), multipotent progenitors (Lin^-^Sca1^+^cKit^+^CD150^-^CD48^-^, MPPs) (**D**), Lin^-^Sca1^+^cKit^+^ CD150^+^CD48^+^ (**E**), and Lin^-^Sca1^+^cKit^+^ CD150^-^CD48^+^ (**F**) in LSK cells of *Zbtb33* knockout mice compared with the wildtype littermates. (**G**) Representative flow cytometry profiles of hematopoietic stem progenitor cells (HSPCs) in bone marrow. (**H**) Representative flow cytometry profiles and quantification of common myeloid progenitors (CMPs) in bone marrow. (**I**) Flow cytometry quantification of T, B cells and neutrophils in bone marrow.

In older *Zbtb33*-KO mice (6-12 months), no statistically significant differences were observed in HSPC subsets or myeloid progenitors compared to aged WT controls (**Supplemental Figure 1C-D**). However, the Lin^-^Sca1^+^cKit^+^CD150^+^CD48^+^ population was significantly reduced in the bone marrow of aged *Zbtb33*-KO mice (**Supplemental Figure 1C**, *P* = 0.0218). Mature cell populations in the bone marrow remained comparable between aged *Zbtb33*-KO and WT mice (**Supplemental Figure 1E**). Overall, *Zbtb33* deficiency was insufficient to promote the expansion of HSPC and progenitors in bone marrow, regardless of age.

### Zbtb33 mutant HSCs have no competitive advantage in competitive transplantation experiments

To assess the hematopoietic reconstitution and expansion potential of *Zbtb33*-deficient HSPCs, competitive bone marrow transplantation assays (cBMT) were performed using a mixture of *Zbtb33* mutant and WT HSCs in recipients. Unlike the findings by *Beauchamp et al.*^4^, our results demonstrated that the loss of *Zbtb33* did not confer a clonal expansion advantage to HSPCs. As illustrated in **Figure 3A**, BMCs from *Zbtb33*-deficient or WT donors (CD45.2LJ) were mixed with the protective competitor BMCs from BoyJ mice (CD45.1LJ) at a 1:1 or 1:5 ratio and transplanted into irradiated recipient mice (CD45.1LJCD45.2LJ). The loss of *Zbtb33* in HSCs did not hinder hematopoiesis reconstruction, as all recipients resumed hematopoiesis and survived throughout the observation period. Peripheral blood chimerism was analyzed by flow cytometry to quantify CD45.2LJ cells. The results indicated that *Zbtb33*-deficient donor cells consistently exhibited lower chimerism compared to WT donor cells, regardless of the donor-to-competitor cell ratio (**Figure 3B-C**).

**Figure 3.**
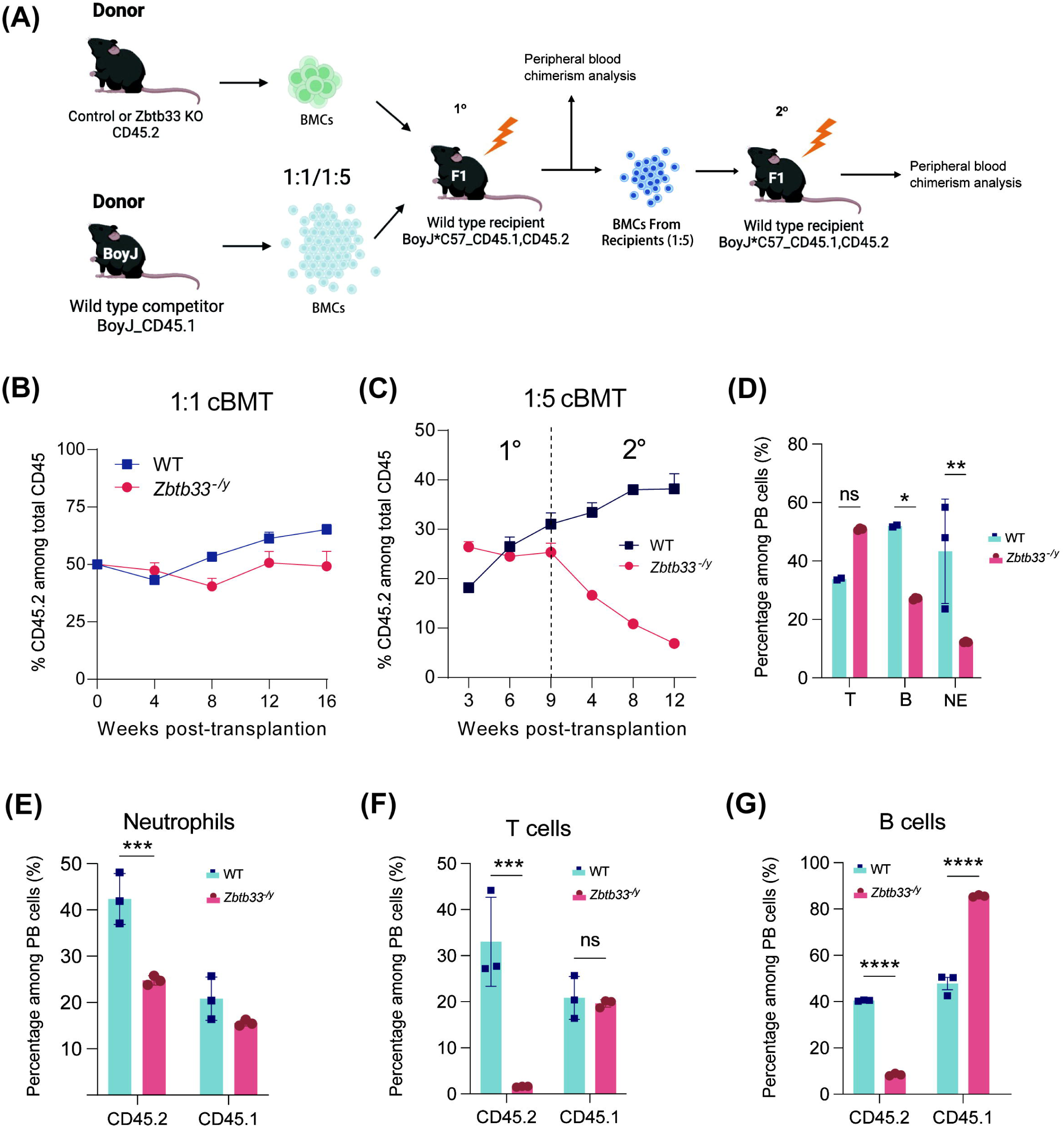
*Zbtb33* mutant HSCs have no competitive advantage in competitive bone marrow transplantation (cBMT) assays. **(A)** Schematic diagram of cBMT assay with a ratio 1:1 or 1:5. **(B)** PB chimerism at the indicated time points with a 1:1 BM cells mixing ratio. **(C)** PB chimerism at the indicated time points post cBMT with a 1:5 BM cells mixing ratio. A secondary cBMT, was performed, with the primary and secondary cBMT separated by the dotted line. **(D)** Frequencies of the indicated cell types in PB at the time point 12 weeks post the cBMT. (**E**-**G**) Flow cytometry quantification of the lineage chimerism analysis of CD45.1^+^ and CD45.2^+^ cells in PB.

In secondary transplantation experiments (1:5 ratio), the decline in CD45.2LJ chimerism was even more pronounced among recipients receiving *Zbtb33*-deficient cells (**Figure 3C**). These results suggested that *Zbtb33*-deficient HSPCs possessed reduced self-renewal capacity (rather than an enhanced capacity) and could not drive clonal hematopoiesis. At the endpoint, significant reductions in neutrophils and B cells of PB were observed in recipients transplanted with *Zbtb33*-deficient BMCs (**Figure 3D**). Further analysis showed no lineage bias, with all peripheral blood cell types exhibiting low chimerism (**Figure 3E-G**).

### Loss of Zbtb33 results in elevated neutrophils of peripheral blood in response to inflammatory stimulus

Under physiological conditions, *Zbtb33* deficiency did not cause significant hematopoietic alterations. To further investigate the hematopoietic responses to external stimuli, we examined the effects of *Zbtb33* deletion under bacterial inflammatory induced by lipopolysaccharide (LPS) and non-bacterial enteritis mediated by dextran sulfate sodium salt (DSS) (**Figure 4A** and **C**).

**Figure 4.**
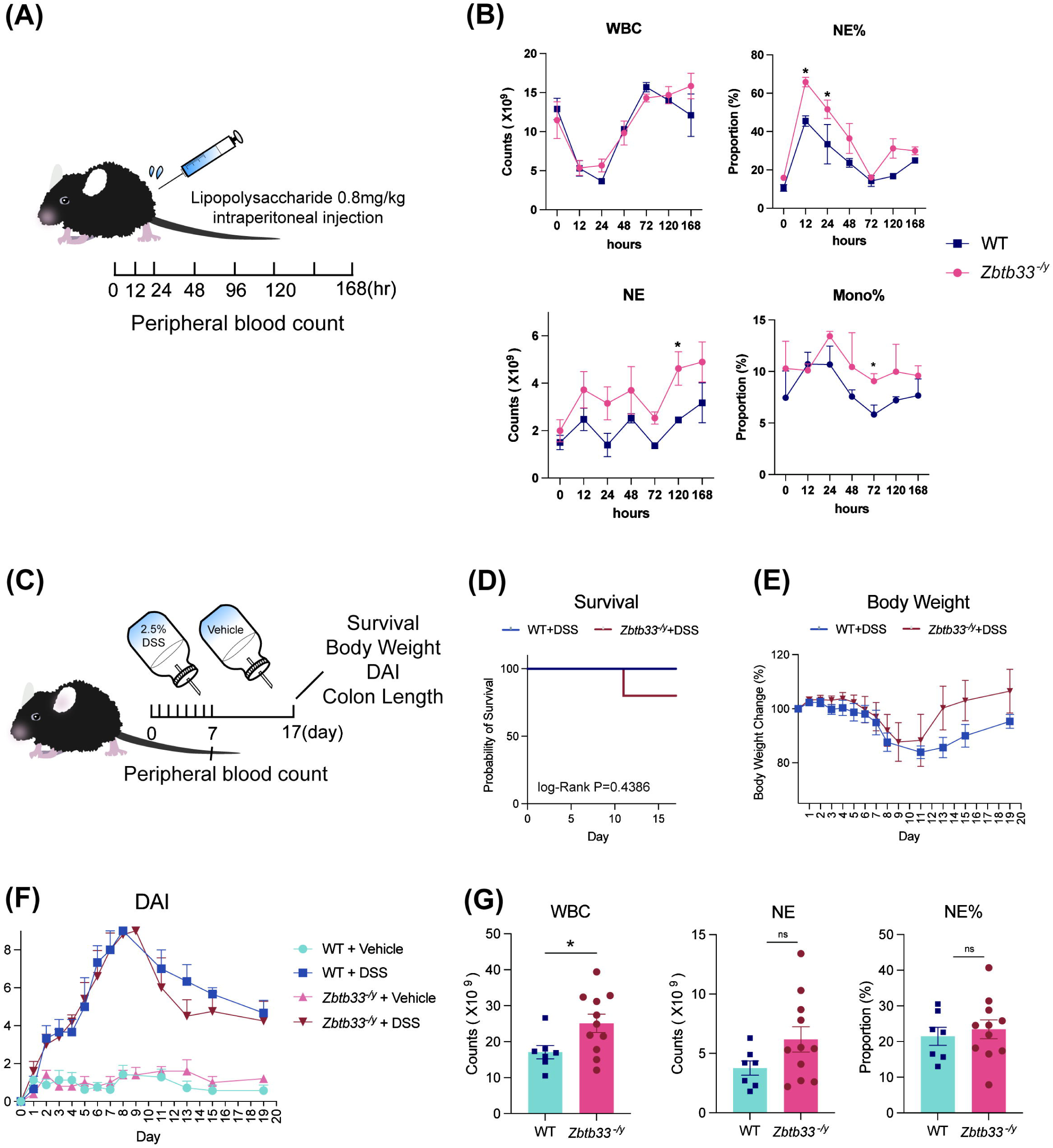
Role of *Zbtb33* upon inflammatory challenges. (**A**) Schematic diagram of an inflammatory model by using lipopolysaccharide (LPS). **(B)** Changes of peripheral blood cells at the indicated time point. **(C)** Schematic diagram of another inflammatory challenge model using Dextran Sulfate Sodium (DSS). **(D)** Survival curves during a DSS-challenge cycle (induction phase and recovery phase). **(E)** Body weight changes during an entire DSS-challenge cycle. **(F)** Disease activity index (DAI) changes during a DSS-challenge cycle. **(G)** Peripheral blood cell counts at 7 days post-DSS feeding.

Upon LPS injection, both *Zbtb33*-knockout and wild-type (WT) mice exhibited a similar trend in white blood cell (WBC) count changes, characterized by an initial decrease (0∼24 hours) followed by an increase (after 24 hours). Neutrophil percentages (NE%) peaked at 12 hours post-injection and declined continuously until 72 hours, reaching similar minimal levels in both groups. However, during the acute inflammatory response phase (12–24 hours), *Zbtb33*-deficient mice displayed a significantly higher NE% than WT mice, indicating a more pronounced inflammatory response (**Figure 4B**, *P* < 0.05). Additionally, absolute neutrophil count (NE) remained elevated in *Zbtb33* KO mice throughout the observation period. During the recovery stage post LPS-administration, NE continued to rise in *Zbtb33*-KO mice, whereas WT group exhibited stable or fluctuating NE levels. These results suggest that *Zbtb33* deficiency may enhance neutrophil differentiation or mobilization in response to inflammatory stimuli (**Figure 4B**, *P*<0.05). Monocyte proportions (Mono%) also remained significantly higher in *Zbtb33*-KO mice at any time point after 24 hours post injection, with the largest difference observed at 72 hours (**Figure 4B**, *P*<0.05). No significant changes were noted in other hematological parameters (**Supplemental Figure 2A**).

For the DSS-induced enteritis model, four groups were rigorously studied: *Zbtb33*-deficient mice and WT mice treated with DSS or vehicle (**Figure 4C**). No significant differences were observed in survival rates, body weight changes, or disease activity index (DAI) scores between *Zbtb33* KO and WT mice with DSS treatment (**Figure 4D-F**). Peripheral blood analysis on the final day of DSS treatment revealed significant higher WBC counts in *Zbtb33*-knockout mice (**Figure 4G**, *P*<0.05), with elevated neutrophil proportions and counts consistent with LPS-induced findings (**Supplemental Figure 2B**). Moreover, colon length is an indicator of colitis severity. After the recovery period following DSS treatment, the colons of *Zbtb33*-KO mice did not demonstrate significant shortening (**Supplemental Figure 2C**; *P*<0.05). Spleen weight did not show significant differences between groups (**Supplemental Figure 2D**).

*Zbtb33* deficiency enhances neutrophil levels during inflammatory responses, with elevated neutrophil counts persisting even during the recovery phase. However, this effect does not significantly impact the overall progression or severity of inflammatory conditions, suggesting that *Zbtb33* deficiency modulates neutrophil dynamics but is insufficient to drive disease outcomes.

### Loss of Zbtb33 did not enhance or promote effects of Tet2 loss-of-function mediated clonal hematopoiesis

CH is a complex consequence of interactions between multiple genetic drivers and environmental factors.^6^ *Zbtb33* as a candidate CH drivers, encodes transcription factor KAISO, which has been reported as a methylated DNA reader^27^. *Tet2*, a common clonal hematopoiesis-associated gene, is closely linked to DNA methylation. Previous studies, including our own, have shown that *Tet2* loss of function (LOF) mutations promote HSPCs self-renewal and mediate myeloid-skewing differentiation. Using the BloodSpot database,^28^ we found that ZBTB33 expression is elevated in early and intermediate stage of myeloid differentiation, such as CMPs, early and late promyelocytes (PMs), while TET2 expression increases during late myeloid differentiation, especially in polymorphonuclear neutrophils (PMNs) (**Supplemental Figure 3A**).

To investigate potential genetic interactions between *Zbtb33* and *Tet2*, we crossed *Zbtb33*-deficient mice with mice harboring the *Tet2* LOF mutations (**Figure5**). Four groups of mice were studied: *Zbtb33* mutant mice (*Zbtb33^-/y^*), *Tet2* LOF heterozygous mutant mice (*Tet2*^+/-^), *Zbtb33* and *Tet2* double mutant mice (*Zbtb33*^-*/y*^; *Tet2*^+/-^), and wild-type littermate controls (WT). *Tet2*-LOF mice exhibited significantly elevated in neutrophils (NE and NE%) and decreased lymphocytes compared to wild-type mice, consistent with the myeloid expansion mediated by *Tet2* mutations. In *Zbtb33^-/y^*; *Tet2^+/-^* mice, we observed elevated WBC, PLT, NE and NE% alongside reduced LY% compared to *Zbtb33^-/y^* mice (**Figure5A**). However, no significant differences were noted between *Tet2^+/-^* and *Zbtb33^-/y^; Tet2^+/-^*mice, indicating that *Zbtb33* mutation had minimal effects on peripheral blood parameters under *Tet2* LOF conditions. Spleen weights showed significant increases in *Zbtb33^-/y^*; *Tet2*^+/-^ mice compared to *Zbtb33^-/y^* littermates, but no differences when compared to the *Tet2^+/-^* mice (**Figure 5B**).

**Figure 5.**
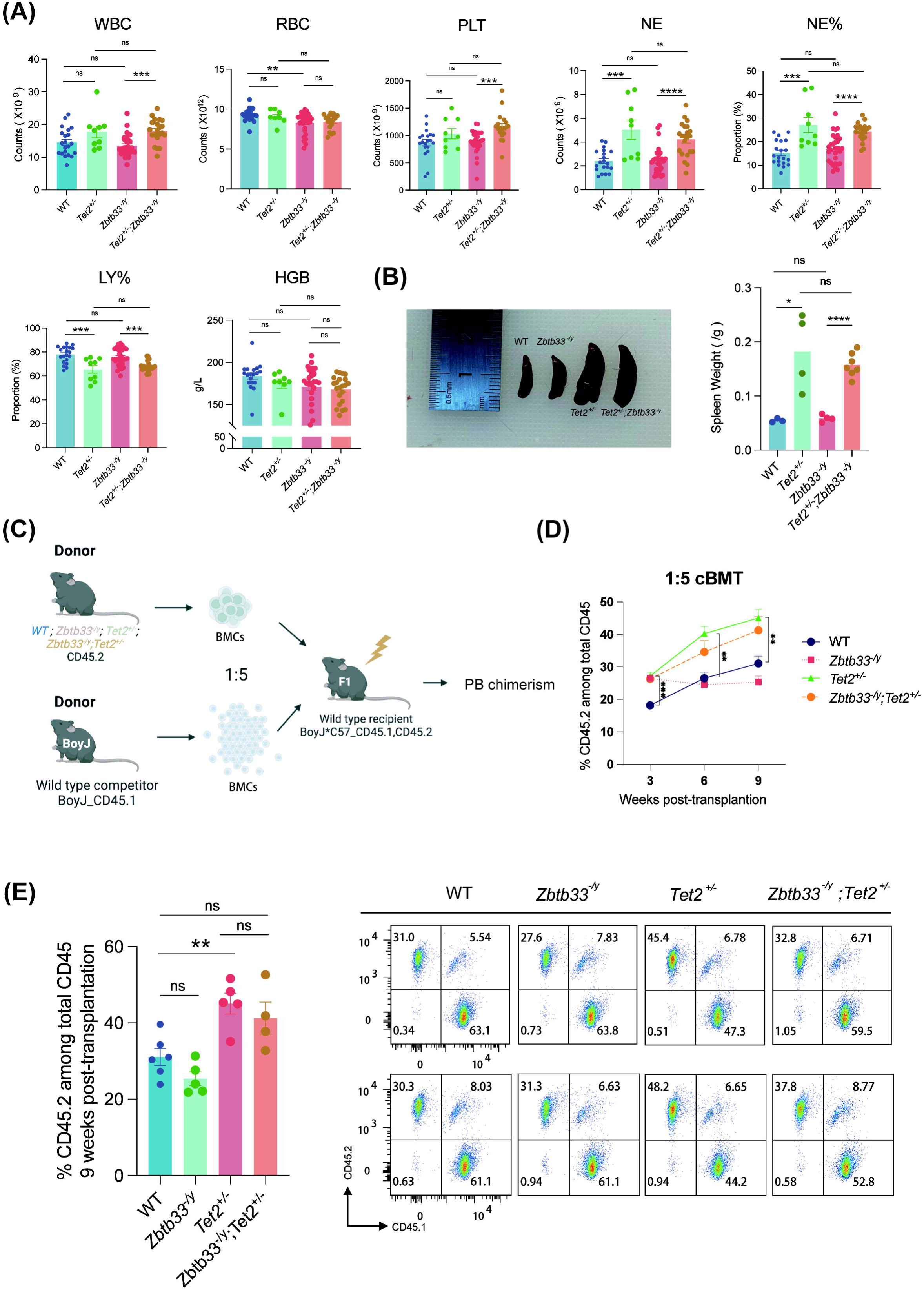
The absence of *Zbtb33* did not enhance or dimmish effects of *Tet2* loss-of-function. **(A)** Blood cell counts in WT, *Tet2^+/-^*, *Zbtb33^-/y^*, and compound mutants *Tet2^+/-^; Zbtb33^-/y^*. **(B)** Representative images and weight measurement of spleens in indicated genotypes. **(C)** Schematic diagram of cBMT assay for in mice with indicated genotypes. Mixing ratio: 1:5. **(D)** PB chimerism analysis at the indicated time points with 4 different genetic background donors. **(E)** Representative flow cytometry profiles and quantification results of PB chimerism at monitoring endpoints (9 weeks post cBMT)

We performed competitive bone marrow transplantations (cBMT) at a 1:5 ratio to assess the clonal expansion of HSPCs with *Zbtb33* and *Tet2* mutations (**Figure5C**). Recipients of *Tet2*^+/-^ BMCs showed significantly higher PB chimerism than WT donors, confirming *Tet2*-LOF mutation-mediated clonal expansion. In contrast, *Zbtb33^-/y^* BMCs exhibited the lowest chimerism. Interestingly, while double mutants BMCs displayed greater clonal expansion than WT or *Zbtb33*^-*/y*^ BMCs, their expansion was moderately reduced compared to *Tet2*^+/-^ BM donors (**Figure5D-E**). These findings suggested that loss of *Zbtb33* did not enhance or promote *Tet2*-driven clonal hematopoiesis.

Considering the established role of Kaiso in DNA damage responses in a *P53*-dependent manner,^29^ we also examined the interplay between *Zbtb33* and *P53*, another key driver of clonal hematopoiesis. Analysis of the BloodSpot database revealed that *P53*, similar to *Zbtb33*, is highly expressed in myeloid lineages of HSPCs, including common myeloid progenitors, granulocyte-macrophage progenitors, and both early and late promyelocytes (**Supplemental Figure 3A**). We generated *Zbtb33*^-/*y*^*; P53^+/-^*mice to assess whether *Zbtb33* deficiency influenced *P53* mutation-driven clonal hematopoiesis. As expected, *P53* mutations caused myeloid expansion, with significant increases in WBC, NE%, and NE. However, no discernible differences in peripheral blood indices were observed between *Zbtb33*^-/*y*^*; P53^+/-^* mice and those with a single *P53* mutation, indicating that *Zbtb33* deficiency dose not significantly alter the hematological changes induced by *P53* mutation (**Supplemental Figure 3B**).

To investigate the impact of *Zbtb33* deficiency on hematopoiesis in an inflammatory context caused by genetic mutations, we crossed *Zbtb33*-KO mice with *Pstpip2* knockout mice, a model that exhibits sterile inflammation^30^. This model mimics the clinical features of chronic multifocal osteomyelitis (CMO) and shows hallmarks of auto-inflammatory disease, including extramedullary hematopoiesis and joint swelling. We conducted a three-month observational study on the four groups of mice: Wild-type (WT) mice, *Zbtb33*-deficient mice (*Zbtb33*^-/*y*^), *Pstpip2* mutant mice (*Pstpip2*^-/*-*^), *Zbtb33* and *Pstpip2* double-mutant mice (*Pstpip2*^-/*-*^;*Zbtb33*^-/*y*^). Through tracking and scoring the number of swollen hind paws over time, we observed a progressive escalation of osteomyelitis symptoms, as indicated by an increasing count of swollen paws with age in both *Pstpip2* mutant groups. Notably, no significant differences in paw swelling were observed between the *Pstpip2* double-mutant and single-mutant mice (**Supplemental Figure 3C**). PB analysis of 3-month-old mutant mice showed a significant increases in neutrophil count and proportion in *Pstpip2*^-/*-*^ mice compared to WT mice. However, no notable differences were observed between double-mutant mice and *Pstpip2* single-mutant mice. These results indicated that *Zbtb33* deficiency did not exert a substantial influence on the myeloid expansion driven by the *Pstpip2* mutation (**Supplemental Figure 3D**).

### Loss of Zbtb33 in tumor microenvironment promoted the tumors growth

To investigate the potential molecular alterations in the hematopoietic system caused by *Zbtb33* deficiency, we performed bulk RNA-seq on bone marrow cells from wild-type mice and *Zbtb33*-deficient mice (n=4 per group). Our differential gene expression analysis identified the top 10 differentially expressed genes in *Zbtb33*-knockout mice (**Figure 6A**). Among the upregulated genes, *Hspb8*, which encodes small heat shock protein B8 stood out. *Hspb8* exhibits dual roles in tumorigenesis, either promoting or suppressing tumor growth depending on the tumor type.^32^ Previous studies have shown that *Hspb8* mediates autophagy in melanoma, thereby inhibiting tumor cell growth and thus exerting anti-tumor effects. The heatmap revealed a significant increase in the expression of *Igkv4* (an immunoglobulin kappa variable gene) in *Zbtb33*-deficient BMCs, suggesting immune dysregulation in the bone marrow. Additionally, *Platr22*, a long non-coding RNA known to influence cell fate and stem cell pluripotency through epigenetic regulation, was markedly upregulated in *Zbtb33*-deficient mice. Another upregulated gene, *Apob*, which encodes apolipoprotein B protein, is involved in the transport of low density lipoprotein (LDL) and very low density lipoprotein (VLDL), and is closely associated with cardiovascular disease.^33^ In contrast, significantly downregulated genes included *Nign1*, *Ren1*, and *Ren2*, which are linked to the ACE inhibitor pathway. These findings suggest that *Zbtb33* deficiency may regulate tumorigenesis, immune responses, cell fate, stem cell pluripotency, and cardiovascular diseases (**Figure 6A**).

**Figure 6.**
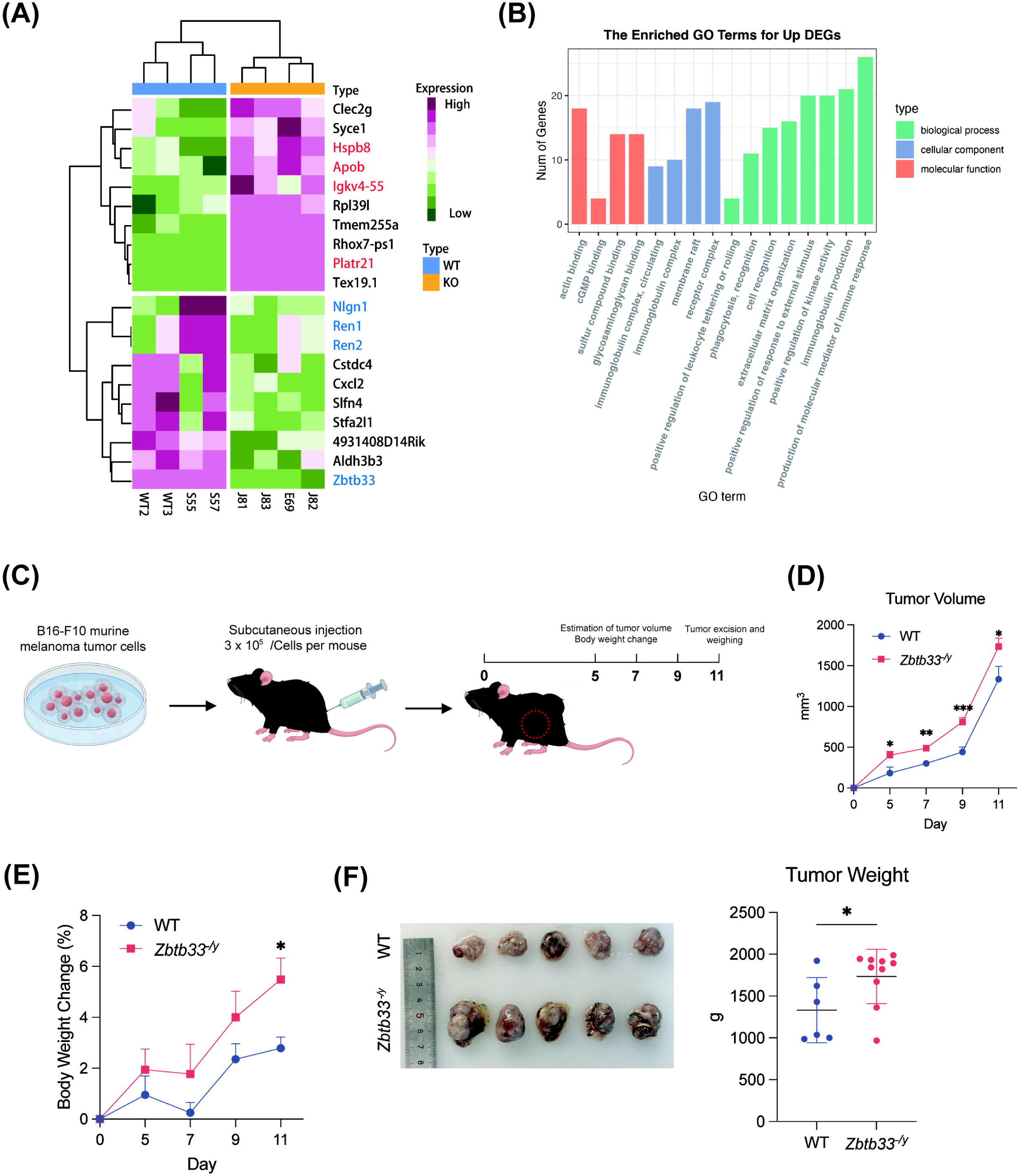
Loss of *Zbtb33* in the tumor microenvironment exacerbates tumor growth. (**A**) RNA-seq analysis of *Zbtb33*-KO bone marrow cells. Heatmap highlights differentially expressed genes (DEGs). The purple grids are for upregulated genes while the green for downregulated genes. (**B**) The top upregulated Gene Ontology (GO) pathways in *Zbtb33*-KO bone marrow cells primarily included immune-related processes. (**C**) Schematic diagram of subcutaneous B16-F10 murine melanoma formation model. (**D**) Tumor volume was calculated. (**E**) Percentage change in body weight of B16-F10 tumor-bearing mice. (**F**) Image and weight of tumors from the indicated host mice.

Gene Ontology (GO) enrichment analysis highlighted upregulated immune response-related pathways in *Zbtb33*-deficient BM cells, including the production of molecular mediators of immune response, immunoglobulin production and circulating, and receptor complex formation (**Figure 6B**). This prompted us to investigate the impact of *Zbtb33* deficiency on the immune microenvironment. We preliminarily tested the role of *Zbtb33* in the context of tumorigenesis by hosting B16-F10 melanoma cells in *Zbtb33*-null mice (**Figure 6C**). Post-subcutaneous injection of B16-F10 cells, we observed a significant exacerbation in tumor size and body weight changes in *Zbtb33*-null mice relative to WT controls (**Figure 6D-F**, *P*<0.05). These findings underscore the potential role of *Zbtb33* as a protective factor within the tumor microenvironment. In conclusion, the preliminary results suggest that absence of *Zbtb33* reshapes immune surveillance, highlighting its underlying function as a tumor suppressor gene.

## Discussion

Clonal hematopoiesis (CH) is defined by age-related expansion of premalignant hematopoietic stem cells (HSCs) with somatic mutations in certain CH-driver genes (i.e. *TET2* and *TP53*). Intriguingly and importantly, this phenomena is associated with the development and progression of various diseases, encompassing both hematological and non-hematological conditions such as Alzheimer’s disease, cardiovascular diseases, chronic inflammation^34^, and hematopoietic malignancies, particularly myeloid neoplasms^35^. Mutations in *DNMT3A, TET2,* and *TP53* (characterized by loss of function) and in *JAK2* (characterized by gain of function), are well-established driver mutations of CH^36^, supported by clinical evidence and experimental validation. Those CH-associated mutations increase the risk of both hematological malignancies and various chronic diseases associated with CH. However, the full spectrum of mutations that drive CH remains not fully elucidated, and there is an urgent need for experimental validation of potential candidate driver mutations. Studies from clinical hematology and mouse genetic models are both important for validating CH drivers, providing a comprehensive understanding of the genetic underpinnings of this condition and its associated diseases.

In 2021, mutations in *ZBTB33/KAISO* were first reported as a novel candidate driver for CH through the integration of DNA-sequencing datasets from TOPMed and ExAC.^4^ By 2024, the year our manuscript is slated for completion, *ZBTB33* has been reaffirmed as a ‘new fitness-inferred driver’ of CH, based on analyses of whole blood somatic mutations and selection from single-cell-derived hematopoietic colonies.^22^ Our work has conducted an in-depth investigation of the mouse homolog of *ZBTB33*, *Zbtb33*, using mouse models. However, our experimental analyses did not substantiate the notion that the loss of *Zbtb33* drives clonal hematopoiesis in the mouse models. Functional analyses, including competitive bone marrow transplantation (cBMT) experiments, indicated that *Zbtb33* deletion did not promote the renewal and expansion associated with inducing clonal hematopoiesis, as tested at mixing ratios of 1:1 or 1:5. *Beauchamp* et al. conducted minimal experimental work, which included CRISPR/Cas9-based gene knocking-out of *Zbtb33* using mouse primary HSCs and transplantation.^4^ The discrepancy in findings is probably due to technical differences.

Our previous studies, supported by corroborating evidence from other research, indicates that the loss of function of the DNA demethylase *Tet2*, a pivotal driver in clonal hematopoiesis (CH), enhances the self-renewal ability of hematopoietic stem cells (HSCs) and predominantly steers their differentiation towards myeloid lineages.^37,38^ Concurrently, mutations in DNA damage repair gene *TP53* represent another prevalent driver category of CH. It was reported that *TP53* has been implicated in regulating Kaiso expression.^29,39,40^ Fascinated by the potential synergistic effects of *Zbtb33* mutations in conjunction with *Tet2* or *Tp53* mutations in CH, we conducted a series of experiments. However, neither the *Zbtb33^-/y^;Tet2^+/-^* nor *Zbtb33^-/y^;Tp53^+/-^* knockout mice models exhibited an accelerated progression of CH compared to their respective *Tet2^+/-^* or *P53^+/-^* single-mutant controls. Similarly, *Zbtb33* deficiency was not sufficient to enhance or alter the phenotypes in well-established chronic osteomyelitis models mediated by *Pstpip2* mutations. This could indicate that the synergy effects of ZBTB33 with TET2, TP53, and PSTPIP2 in hematopoiesis are limited, and the impact of ZBTB33 in clonal hematopoiesis is comparatively marginal when contrasted with that of other well-characterized driver genes.

Under physiological conditions, *Zbtb33^-/y^* mice did not exhibit abnormalities. However, we are intrigued by the possibility that these mutant mice might exhibit aberrant phenotypes when confronted with pathogens or inflammation. Previous studies have indicated that overexpression of KAISO, which encodes by *ZBTB33*, can promote intestinal inflammation, which is associated with altered p120ctn function and potentially linked to the disruption of the intestinal immune barrier and increased intestinal permeability.^41^ In our study, we assessed the responses of *Zbtb33*-null mice to lipopolysaccharide (LPS) and dextran sulfate sodium (DSS) challenges. In our experimental setup, these two different external inflammation stimuli did not affect mouse survival. Moreover, upon LPS challenge or DSS treatment, the phenotypic indicators suggested that the phenotypes of *Zbtb33* mutants were largely indistinguishable from those of wild-type (WT) controls, with the exception of a slight increase in neutrophils percentages (NE%, **Figure 4**) in peripheral blood of *Zbtb33^-/y^* mice. These findings suggested a minimal or dispensable role of *Zbtb33* in inflammation resolution.

In addition, the role of *Kaiso* in oncogenesis has indeed been a subject of controversy,^42^ with evidence suggesting its dual capacity to act as both transcription activator and repressor.^43^ Identified as a tumor suppressor,^44^ Kaiso’s repression of key target genes such as Matrilysin, Siamois, and Wnt11 has been implicated as a crucial mechanism in its antitumorigenic activity.^6,7,8^ Our findings has revealed that the deletion of Kaiso in the immune environment can paradoxically accelerate tumor growth in a murine B16-F10 melanoma tumor model, which indicated that the absence of *Zbtb33* may reshape immune regulation and function as a tumor suppressor gene. As we provided preliminary evidence suggesting loss of ZBTB33/Kaiso impaired the antitumor immune response using a single mouse cell line, further experimental validation using a broader range of mouse tumor cell lines and in-depth mechanistic exploration through scRNA-seq analysis is demanded.^47^

In conclusion, the study provides experimental results suggest that: 1) Similar to *Tet2* and *Tp53*, *Zbtb33/Kaiso* is broadly expressed across various hematopoietic cell types; 2) Functionally, we have demonstrated that *Zbtb33^-/y^* mutant mice (male) appear physiologically normal and do not exhibit any overt abnormalities under normal conditions; 3) The loss of *Zbtb33* in HSCs did not promote clonal hematopoiesis; 4) When challenged with LPS or DSS, or when bred with *Pstpip2^-/-^* mice, *Zbtb33^-/y^*mice failed to manifest abnormalities in inflammation resolution; and 5) Interestingly, *Zbtb33^-/y^* mice transplanted with murine tumor cells showed increased tumor growth, suggesting an anti-tumor function of *Zbtb33* in the tumor microenvironment. Given that our findings are based solely on the *Zbtb33-*KO mouse model, further experimental analysis using human bone marrow cells is necessary to determine the inclusion or exclusion of ZBTB33 in the gene-mutation panel tests for clonal hematopoiesis (CH) and myelodysplastic syndromes (MDS). This additional research will help to elucidate the role of ZBTB33 in human disease contexts and guide clinical decision-making.

### Conclusion

This study shows that *Zbtb33* deficiency does not drive clonal hematopoiesis or enhance the expansion of hematopoietic stem cells, even in combination with *Tet2* or *Tp53* mutations. While *Zbtb33* does not play a direct role in CH, it may regulate the tumor microenvironment and influence cancer immunity. These findings suggest that ZBTB33 plays a limited role in hematopoiesis, with its involvement in immune regulation requiring further investigation.

## Acknowledgments

We thank our colleagues for technical support, critically reading our manuscript, and their suggestions to improve the manuscript. We would also like to thank Dr. Hu for sharing us mouse tumor cell lines.

## Authorship

Conceptualization: ZC

Methodology: YL, KL, JL

Investigation: YL, KL, JL

Visualization: YL, KL

Supervision: ZC, ZZ, GD

Writing—original draft: YL, KL, ZC

Writing—review & editing: ZC, ZZ

## Conflict of interest disclosure

ZC is a scientific advisor to Beijing SeekGene BioSciences Co. Ltd. The other authors declare no potential conflict of interest.

## Funding

This work was supported in part by grants from the Tianjin Medical University Talent Program, Tianjin Municipal Education Commission Scientific Research Program Projects (#2022KJ194 to GD), and The National Natural Science Foundation of China (82170173 and 82371789 to ZC).

## Methods

### Generation of Zbtb33 deficient transgenic mice

CRISPR/Cas9 technology was employed to target the *Zbtb33* gene located on the X chromosome of C57BL/6 mice for knockout. A specific sgRNA was designed, and high-throughput electroporation of fertilized eggs was conducted to generate Zbtb33 gene knockout mice, with assistance from Cyagen US Inc. C57BL/6 (CD45.2^+^) and BoyJ (CD45.1^+^) mice were purchased from Cyagen US Inc. and used in the experiments. All animals were housed under specific pathogen-free conditions with a 12-hour light/dark cycle at Tianjin Medical University. The study included younger adult mice aged 8 weeks and older mice aged 6∼12 months. The animal protocols were provided by Cyagen US Inc., and the experiment was conducted in accordance with the regulations of the Animal Ethics Committee of Tianjin Medical University, with their approval.

### Genotyping and PCR

Transgenic pups were identified through polymerase chain reaction (PCR) analysis of DNA extracted from tail biopsies. The primer pairs used specific to sequences within the murine *Zbtb33* gene:

Forward Primer (F1): 5’-TATGGGTCCTTTGCACTTCAGAT-3’
Reverse Primer (R1): 5’-CACTGAAAACTTCATCTTCACACCT-3’

The PCR was conducted following standard protocols.

### Complete blood count (CBC) procedure

Blood was collected from the outer canthus of mice using a capillary glass tube (Kimble) and transferred into an EDTA-containing EP tube. The EDTA and blood were thoroughly mixed for 30 seconds using a mixing device (Kylin-Bell). Subsequently, the proportions of blood cells were determined using an automated complete blood count machine (SHENGCHENYILIAO).

### Mouse tissue collection

Mice were euthanized by CO_2_ asphyxiation, and the abdominal cavity was exposed. Spleens, colons, and tumors were removed and immediately placed in pre-cooled phosphate-buffered saline (PBS) (Solarbio) on ice for subsequent investigation. Bone marrow cells were harvested from the femurs and tibias of mice by flushing the bone marrow cavity with sterile PBS. The cell suspension was filtered through a 70 μm cell strainer (BD Biosciences) and washed twice with PBS.

### Flow cytometry analysis of peripheral blood cells and bone marrow cells

Purified murine peripheral blood mononuclear cells (PBMCs) were stained with the following surface marker: CD3-PE, CD19-APC, GR1-PE, Mac-1/CD11b-FITC (all from BD Biosciences). The hematopoiesis stem progenitor cells subpopulations were characterized with the following staining strategies: Lin-PE, cKit-APC, Sca1-APC-Cy7, CD150-PE-Cy5.5, CD48-PE-Cy7 (all from BD Biosciences). For myeloid progenitor cell subsets, the following antibodies were used: cKit-APC, Sca1-APC-Cy7, CD34-FITC, CD16-PE-Cy7 (BD Biosciences). Flow cytometry was performed utilizing a BD LSRFortessa analyzer, and data were analyzed using FlowJo software (version 10.8.1, FlowJo LLC). Gating strategies were established based on forward and side scatter to remove debris and doublets, and specific cell populations were identified based on their surface marker.

### Competitive bone marrow transplantation

Bone marrow cells (BMCs) were harvested from either wild-type or *Zbtb33* knockout mice with a C57BL/6 background (CD45.2^+^). First-generation (F1) mice (CD45.1^+^; CD45.2^+^) were generated by crossing C57BL/6 wild-type mice (CD45.2^+^) with BoyJ mice (CD45.1^+^). F1 recipient mice underwent lethal irradiation with doses of 10 Gy and 7 Gy, separated by a 4-hour interval. BMCs from wild-type or *Zbtb33* knockout mice (CD45.2^+^) were mixed with BoyJ competitor cells (CD45.1^+^) at ratios of 1:1 and 1:5 and were injected into the tail vein of F1 (CD45.1^+^; CD45.2^+^) mice 24 hours post-irradiation. The chimerism of CD45.2^+^ cells in the peripheral blood of recipient mice was evaluated every 3 or 4 weeks during the transplantation period. For secondary transplantations, bone marrow cells derived from the first recipients were transplanted into each lethally irradiated recipient. Peripheral blood chimerism was measured every 4 weeks, and bone marrow chimerism was observed at the endpoint of the observation period.

In additional competitive bone marrow transplantation experiments to assess the effects of *Zbtb33* on hematopoiesis in *Tet2*-mutant mice, donors included *Zbtb33^-/y^/Tet2^+/-^*single and double mutant mice and wild-type mice, with F1 mice serving as recipients.

### Mouse inflammatory stimulus experiment

Lipopolysaccharide (LPS), derived from Escherichia coli (Sigma, L2880) was administered intravenously to mice at a dose of 0.8 mg/kg. The control group received an equal volume of phosphate-buffered saline (PBS) as the vehicle. Peripheral blood samples from these mice were collected and sequentially monitored at 12, 24, 48, 96, 120, 168 hours post-administration.

Dextran sulfate sodium salt (DSS) was obtained from MP Biomedicals and dissolved in sterile water to prepare a 2.5% solution. Mice were randomly assigned to four groups: *Zbtb33* mutant mice fed with DSS (*Zbtb33^-/y^* + DSS), wild-type mice fed with DSS (WT + DSS), *Zbtb33* mutant mice fed with vehicle (*Zbtb33^-/y^* + vehicle), and wild-type mice fed with vehicle (WT + vehicle). For the initial 7 days, the experimental groups (WT + DSS; *Zbtb33^-/y^* + DSS) were fed 2.5% DSS to induce non-bacterial colitis, followed by a 10-day recovery phase during which they were fed with the vehicle solution. Meanwhile, the control groups (WT + vehicle; *Zbtb33^-/y^* + vehicle) were fed the vehicle solution throughout the entire period. After completing the entire cycle, the survival rate, body weight changes, disease activity index (DAI), and colon length were measured and recorded for each group.

### Disease activity index (DAI) for enteritis model

The Disease Activity Index (DAI) is an important indicator utilized for evaluating the severity of colitis. DAI scores include factors such as changes in body weight, stool consistency, and the presence of occult blood in stools^48,49^. The fecal occult blood test was performed using an o-toluidine-based test kit, along with its instruction manual, provided by Beijing Leagene Biotechnology Co., Ltd. In the o-toluidine-based fecal occult blood test, the reagent reacts chemically with blood in the stool, causing a color change, typically to blue or blue-green, when blood is present. The speed and intensity of the color change indicate the severity of occult bleeding (Table 1).

**Table 1.**
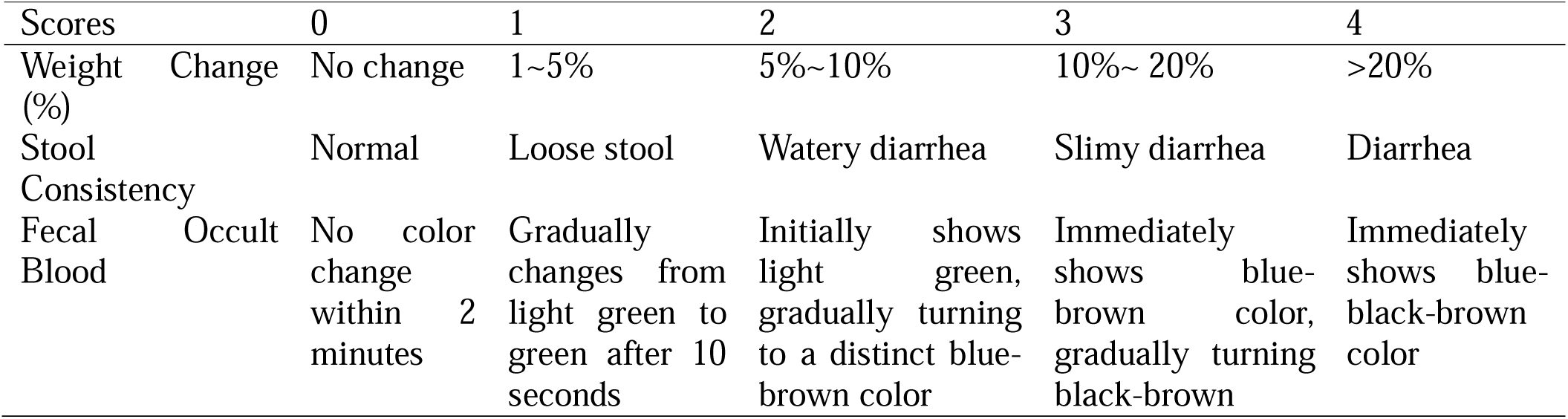
Evaluation indicators of DAI.

### Subcutaneous tumor formation model

B16-F10 melanoma tumor cells were provided by Professor Deqing Hu from the School of Basic Medicine at Tianjin Medical University. The cells were stored in liquid nitrogen and cultured in RPMI1640 medium (Meilunbio) supplemented with 10% fetal bovine serum (ExCell Bio). The cells were maintained under standard conditions until they reached logarithmic growth. The collected cell suspension was centrifuged at 1000 rpm for 5 minutes and washed twice with PBS. The cells were then resuspended in PBS to a final concentration of 6 × 10^6 cells/mL. The cell suspension was mixed with Matrigel at a 1:1 ratio in a 4°C environment to prepare a final concentration of 3 × 10^5 cells/mL. Prior to inoculation, mice were weighed to ensure their weight was between 18 and 22 grams. The hair on the lower right side of their back was shaved using a mouse hair trimmer to expose the skin, which was then disinfected with 75% ethanol. The cell suspension was gently resuspended and was slowly injected subcutaneously into the mouse with at a volume of 100 µL.

Tumor growth was monitored by measuring the maximum diameter (L) and minimum diameter (W) of the tumors every two days with a vernier caliper. Tumor volume (V) was calculated using the formula (V = L * W^2^/2, L= maximum diameter of tumor, W= minimum diameter of tumor).

When the tumor volume reached approximately 2000mm^3^, all mice were humanely euthanized by exposure to carbon dioxide.

### Bulk RNA sequencing of bone marrow cells

Bone marrow cells were obtained from *Zbtb33^-/y^* mice and their wild-type littermates. RNA was isolated and extracted using the RNeasy mini kit (Qiagen) following the manufacturer’s instructions. The quality and quantity of the RNA were assessed using a NanoDrop spectrophotometer (Thermo Fisher Scientific) and an Agilent 2100 Bioanalyzer (Agilent Technologies). Bulk RNA-seq was performed at Majorbio using the NovaSeq 6000 platform (Illumina). Libraries were prepared using the TruSeq RNA Sample Preparation Kit (Illumina) according to the standard protocol. The resulting libraries were sequenced on the NovaSeq 6000 platform to generate 150 bp paired-end reads.

Raw sequencing reads were processed using the fastp tool (version 0.20.0) to perform quality control and trimming. The cleaned reads were aligned to the reference genome (Mus musculus, GRCm38) using the STAR aligner (version 2.7.3a). Aligned reads were quantified using featureCounts (version 1.6.4) from the Subread package. The output from featureCounts was used for downstream differential expression analysis.

### Differential expression gene analysis and enrichment analysis

Differential expression gene analyses between *Zbtb33* knockout and wild-type mice (4 mice per group) were performed using R software (version 4.2.1), R Studio, and DESeq2 (version 1.36.0). Genes with *P* values below 0.05 were considered significantly differentially expressed. Gene ontology enrichment analysis was conducted using the clusterProfiler based on log2 fold change results obtained from the differential expression analysis and visualized by ggplot2 R package.^50^

### Data Availability Statement

The scRNA-seq data for this study are available in the Sequence Read Archive (SRA) at NCBI, under the accession number SUB15022287 (https://www.ncbi.nlm.nih.gov/sra). All other relevant data are available upon request from the corresponding authors.

### Statistical analysis

Statistical differences across multiple groups were assessed using a two-way ANOVA with Newmane-Keuls multiple comparison Test, unless specified otherwise. Single comparisons relied on unpaired, two-tailed Student’s t-tests. For survival analysis, Log-rank tests were applied. In general, *P* values < 0.05 were considered significant and were indicated as follows: \**P* < 0.05, ** *P* < 0.005, *** *P* < 0.0005, ns: not significant. Calculations of *P* values were performed using GraphPad Prism 9.0 software.

---

**Supplemental Figure 1, related to Figure 1:**

**(A)** Overview of the targeting strategy for generating *Zbtb33* knockout mice. Exon2 is removed in the *Zbtb33*^-/y^ mice.
**(B)** Bar chart showing various parameters in whole blood counts in aged *Zbtb33*-deficient mice and matched wildtype littermates.
(**C**-**E**) Percentages of indicated hematopoietic stem/progenitor cells and mature cells in bone marrow from aged *Zbtb33* knockout and wildtype mice.

**Supplemental Figure 2, related to Figure 4:**

**(A)** Changes of peripheral blood cells at the indicated time point.
**(B)** Whole blood counts at 7 days post-DSS feeding in the context of loss of *Zbtb33* and wildtype genotype.
**(C)** Representative images and length measurement of integrate colons in DSS-administrated *Zbtb33* knockout mice and the wildtype littermates.
**(D)** Photography and weight of spleens in DSS-induced inflammation models in the context of loss of *Zbtb33* and wildtype genotype.

**Supplemental Figure 3, related to Figure 5:**

**(A)** Expression level of *ZBTB33*, *TET2*, and *TP53* (mRNA, log_2_) in indicated cells (source: the bloodspot database)
**(B)** Blood cell counts in WT, *Tp53^+/-^*, *Zbtb33^-/y^*, and *P53^+/-^; Zbtb33^-/y^*.
(**C**-**D**) Disease score quantification and PB cell counts in WT, *Pstpip2^-/-^*, *Zbtb33^-/y^*, and *Pstpip2^-/-^; Zbtb33^-/y^*.

## Abbreviation

AML: (acute myeloid leukemia)
AS: (Alternative Splice)
BM: (bone marrow)
BMCs: (bone marrow cells)
CBC: (complete blood count)
cBMT: (competitive bone marrow transplantation)
CH: (clonal hematopoiesis)
CHIP: (clonal hematopoiesis of indeterminate potential)
CMO: (chronic multifocal osteomyelitis)
CMPs: (common myeloid progenitors)
DAI: (disease activity index)
DSS: (dextran sulfate sodium salt)
GMPs: (granulocyte-macrophage progenitors)
GO: (gene ontology)
HSCs: (hematopoietic stem cells)
HSPC: (hematopoietic stem and progenitor cell)
HSPCs: (hematopoietic stem progenitor cells)
LDL: (low density lipoprotein)
LOF: (loss of function)
LPS: (lipopolysaccharide)
LY%: (lymphocyte proportion)
MDS: (myelodysplastic syndromes)
Mono%: (monocytes cell proportions)
MPs: (myeloid progenitors)
MPNs: (myeloproliferative neoplasms)
MPPs: (multipotent progenitors)
NE: (absolute neutrophil count)
NE%: (neutrophil proportion)
ns: (not significant)
PB: (peripheral blood)
PBS: (phosphate-buffered saline)
PCR: (polymerase chain reaction)
PLT: (platelet)
PMNs: (polymorphonuclear neutrophils)
PMs: (promyelocytes)
*Pstpip2*: (proline-serine-threonine phosphatase–interacting protein 2)
RBC: (red blood cell)
VAF: (variant allele fractions)
VLDL: (very low density lipoprotein)
WBC: (white blood cell)
WT: (wild-type)
*Zbtb33*: (Zinc Finger and Btb Domain Containing 33)
*Zbtb33*-KO: (*Zbtb33*-knockout mice)

## Notes

https://www.ncbi.nlm.nih.gov/sra

### Competing Interest Statement

The authors have declared no competing interest.

