## Supplementary figures and images for "Loss of transcriptional factor *Zbtb33* fails to induce clonal hematopoiesis in mice but plays a role in tumor immunity"

### Supplementary Figure 1

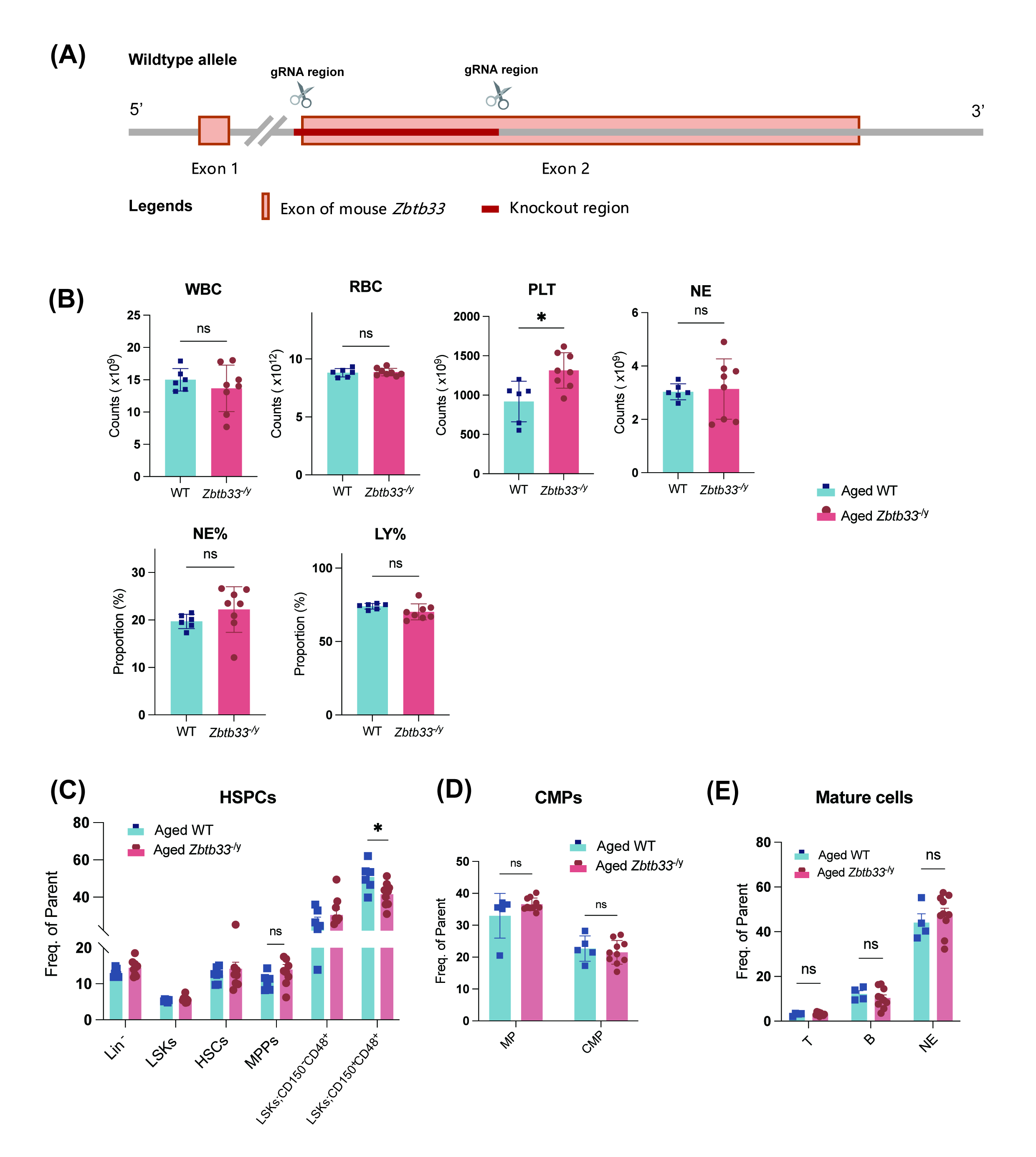

### Supplementary Figure 2

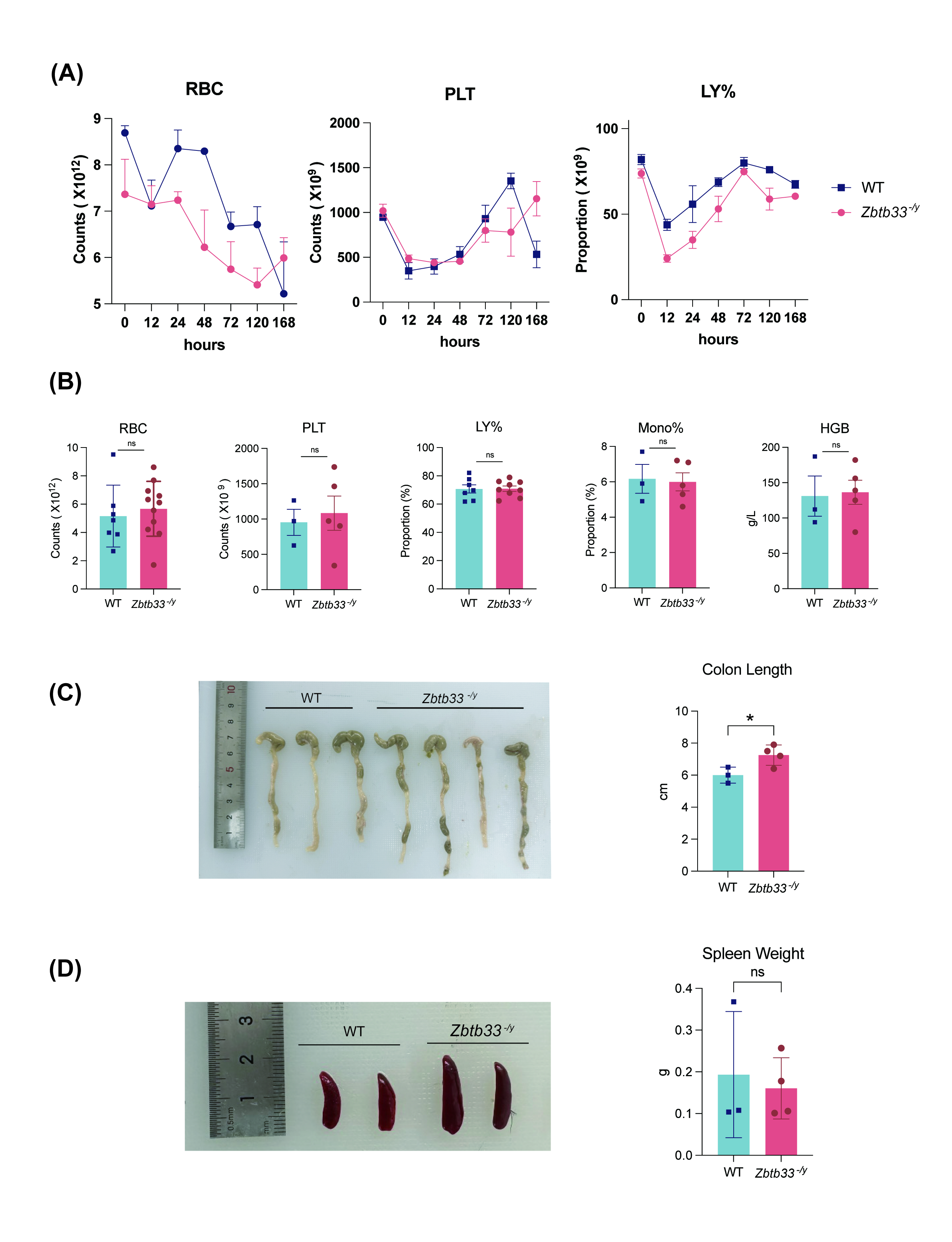

### Supplementary Figure 3

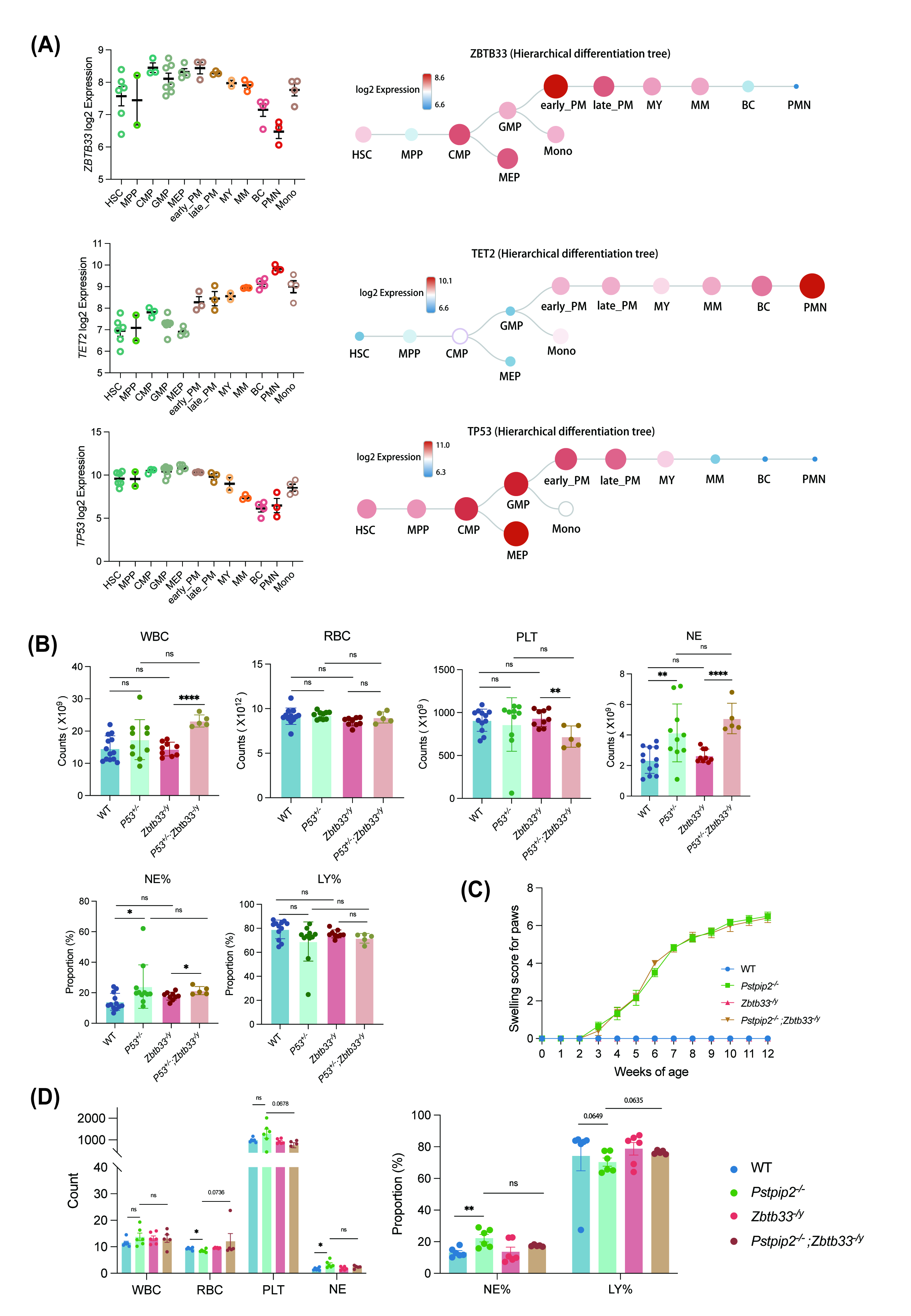
